# The epoxyeicosatrienoic pathway is intact in endothelial and smooth muscle cells exposed to aldosterone excess

**DOI:** 10.1101/2021.02.04.429624

**Authors:** Laura Brunnenkant, Yao Meng, Jing Sun, Jair Gonzalez Marques, Berthold Koletzko, Michael Mederos Y Schnitzler, Thomas Gudermann, Tracy Ann Williams, Felix Beuschlein, Daniel A Heinrich, Christian Adolf, Martin Reincke, Holger Schneider

## Abstract

**Objectives:** Endothelial dysfunction (ED) is considered to be a major driver of the increased incidence of cardiovascular disease in primary aldosteronism (PA). Whether the epoxyeicosatrienoic acid (EET) pathway, involving the release of beneficial endothelium-derived lipid mediators, contributes to ED in PA is unknown. Preclinical evidence suggests this pathway to be relevant in the pathogenesis in various models of experimental hypertension. In addition, an orally available soluble epoxide hydrolase inhibitor, which halts the breakdown of EETs, has already passed a phase 1 clinical trial.

We, therefore, exposed primary human coronary artery endothelial cells to 1 nM aldosterone.

**Methods:** We used qPCR to investigate changes in the expression levels of essential genes for the synthesis and degradation of EETs as well as mass spectrometry to determine endothelial synthetic capacity to release EETs upon stimulation. We also assessed primary human coronary artery smooth muscle cells for expression of putative EET receptor ion channels or downstream mediators as well as for the calcium response to EETs using calcium imaging.

**Results:** No major aldosterone-related expression changes were detected on the endothelial as well as the smooth muscle side. Stimulated release of endothelial EETs was unaffected. Likewise, the smooth muscle calcium response was unchanged after aldosterone excess.

**Conclusions:** The EET pathway is not negatively affected by increased aldosterone concentrations as seen in PA. Modulating the EET pathway with therapeutic intent in patients with PA might therefore be assessed in future preclinical and clinical trials to address ED.

## Introduction

Primary aldosteronism (PA) is the most common cause of endocrine hypertension. In recent years, it has been appreciated that in up to 12 % of patients arterial hypertension can be attributed to aldosterone excess[1]. It is of particular importance to address this condition adequately as high levels of aldosterone cause a significantly greater prevalence of atherosclerotic sequelae as reflected by blood-pressure-independent detrimental effects on the cardiovascular system[2]. In particular, renal injury, atrial fibrillation, coronary artery disease and stroke even when compared to essentially hypertensive control patients who had been matched for age, sex and blood pressure are more abundant in patients with PA. One of the earliest steps in the pathogenesis of atherosclerosis is endothelial dysfunction[3]. In line with previous observations indicating a high prevalence of atherosclerotic disease, endothelial dysfunction has been demonstrated in patients with PA[4–6]. Endothelial dysfunction is a descriptive term which does not allow for a mechanistic attribution. While experimental studies have linked aldosterone-mediated endothelial dysfunction to a release of constrictive prostanoids[7] or endothelin-1[8], most lines of explanation focus on impaired endothelial nitric oxide (NO) generation, decreased NO bioavailability or decreased smooth muscle response to NO[9–11].

However, the relative importance of NO as vasodilator decreases with ageing and sedentary life style[12] and, consequently, a shift to other endothelial mediators seems to occur. Other non-nitric oxide endothelial factors encompass substances like prostacyclin (PGI_2_) and a heterogeneous group of mediators summarized as EDHF (endothelium-derived hyperpolarizing factor). Diverse mechanisms such as localized increases in extracellular potassium via release from the endothelial cytosol, hyperpolarizing currents through myoendothelial gap junctions, release of hydrogen peroxide and arachidonic-acid-derived epoxyeicosatrienoic acids (EETs) have been proposed to constitute molecular correlates of EDHF[13].

Specifically, the role of EETs in aldosterone-mediated endothelial dysfunction has barely been investigated so far. EETs comprise 4 bioactive regioisomers which are named according to the position of the epoxide group (5,6, 8,9, 11,12 and 14,15 EET). They are degraded into inactive DHETs (dihydroxyeicosatrienoic acids) by epoxide hydrolases. Clinical data indicate that levels of circulating 14,15 DHET, the breakdown product of 14,15 EET, are increased in patients with PA, which might point to increased breakdown of active EETs[14]. Further, experimental hypertension induced by administration of the mineralocorticoid precursor DOCA (deoxycorticosterone acetate) was reversed after administration of an inhibitor of soluble epoxide hydrolase[15], which in turn would increase the bioavailability of endothelium-derived EETs. These findings suggest a particular role of EETs in the pathogenesis of mineralocorticoid-induced endothelial dysfunction and warrant further investigation.

Since the impact of aldosterone excess on the EET pathway in both major vascular cell types is unknown, we set out to explore for putative pathologic effects using primary human coronary artery cell lines. We addressed both the capacity of endothelial cells to synthesize and release EETs as well as the calcium response of smooth muscle cells to investigate which cell type might be specifically perturbed. We deliberately focussed on pharmacologic strategies to allow for extrapolation of our results to clinical cohorts.

## Materials and Methods

The authors refer to the online-available supplement with a detailed list of materials and methods used in the experiments.

### Statistics

GraphPad Prism 8 (GraphPad Software, San Diego, CA, USA) was used for construction of graphs and computation of statistical significances. Bar graphs show mean ± SEM. Statistical tests comprised t-tests or 1-Way ANOVA followed by indicated post hoc tests, wherever applicable. Statistical significance levels were set to a two-sided p-value of <0.05.

## Results

In endothelial cells (EC), EETS are synthesized by CYP2C isoenzymes[16,17] as well as CYP2J2[18]. Assessment of the expression status of CYP-epoxygenases revealed that human coronary artery EC express CYP2C8 and CYP2J2. CYP2C9 mRNA could not be detected. Neither CYP2J2 nor CYP2C8 showed significant changes in transcript levels upon challenge with aldosterone, with or without co-stimulation by cortisol (Figure 1A, 1B).

**Figure 1:**
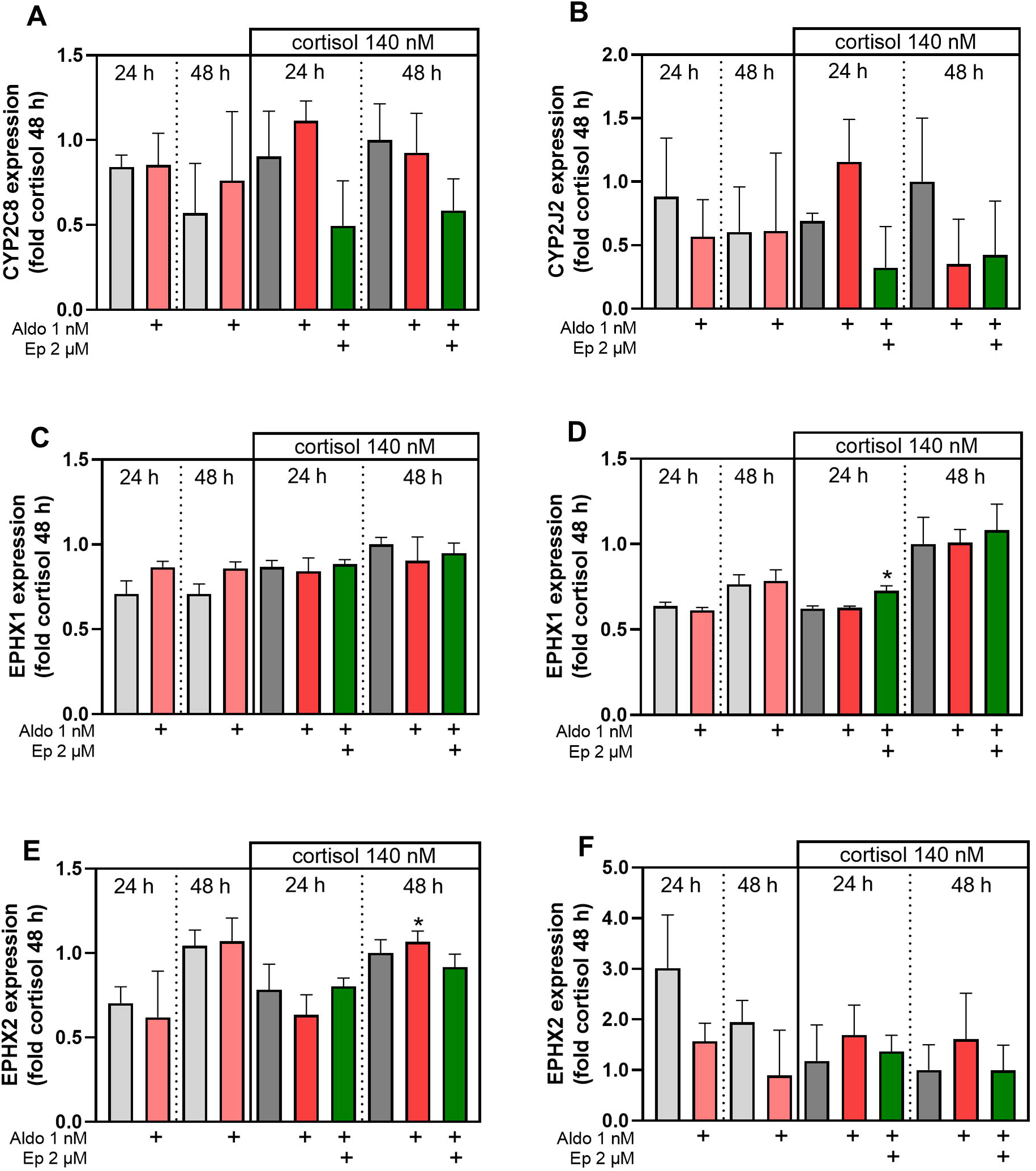
**A, B:** Effect of duration of exposure to 1 nM of aldosterone and co-stimulation with cortisol on CYP2C8 (**A**) and CYP2J2 (**B**) mRNA expression in human coronary artery endothelial cells. Each individual relative expression value was normalized to the mean of the 48 h cortisol control. Aldo, aldosterone, Ep, eplerenone. **C, D:** Effect of duration of exposure to 1 nM of aldosterone and co-stimulation with cortisol on EPHX1 (microsomal epoxide hydrolase) mRNA expression in human coronary artery smooth muscle cells (**C**) and endothelial cells (**D**). Aldo, aldosterone, Ep, eplerenone.*: p<0.05 vs. Aldo+cortisol 24 h, t-test. **E, F:** Effect of duration of exposure to 1 nM of aldosterone and co-stimulation with cortisol on EPHX2 (soluble epoxide hydrolase) mRNA expression in human coronary artery smooth muscle cells (**E**) and endothelial cells (**F**). Aldo, aldosterone, Ep, eplerenone.* p<0.05 vs. Aldo+cortisol 24 h, t-test.

There are at least two epoxide hydrolases which have been reported to be capable of converting EETs to inactive DHETs, microsomal epoxide hydrolase (EPHX1) and soluble epoxide hydrolase (EPHX2), with the latter reportedly being the most important one[19].

Microsomal epoxide hydrolase (EPHX1) and soluble epoxide hydrolase (sEH, EPHX2) were not dysregulated in aldosterone-treated smooth muscle cells (SMC) (Figure 2C, 2E) and EC (Figure 2D, 2F) when compared to their appropriate time-controls. A striking observation, however, was the drastically increased expression of EPHX1 over EPHX2 within both cell types: In 48 h cortisol-treated SMC normalized expression levels (2^-ΔCT^) of EPHX1 versus EPHX2 were 0.24 ± 0.02 versus 0.002 ± 0.0001; 48 h F-treated EC: 0.83 ± 0.13 versus 0.003 ± 0.001.

**Figure 2:**
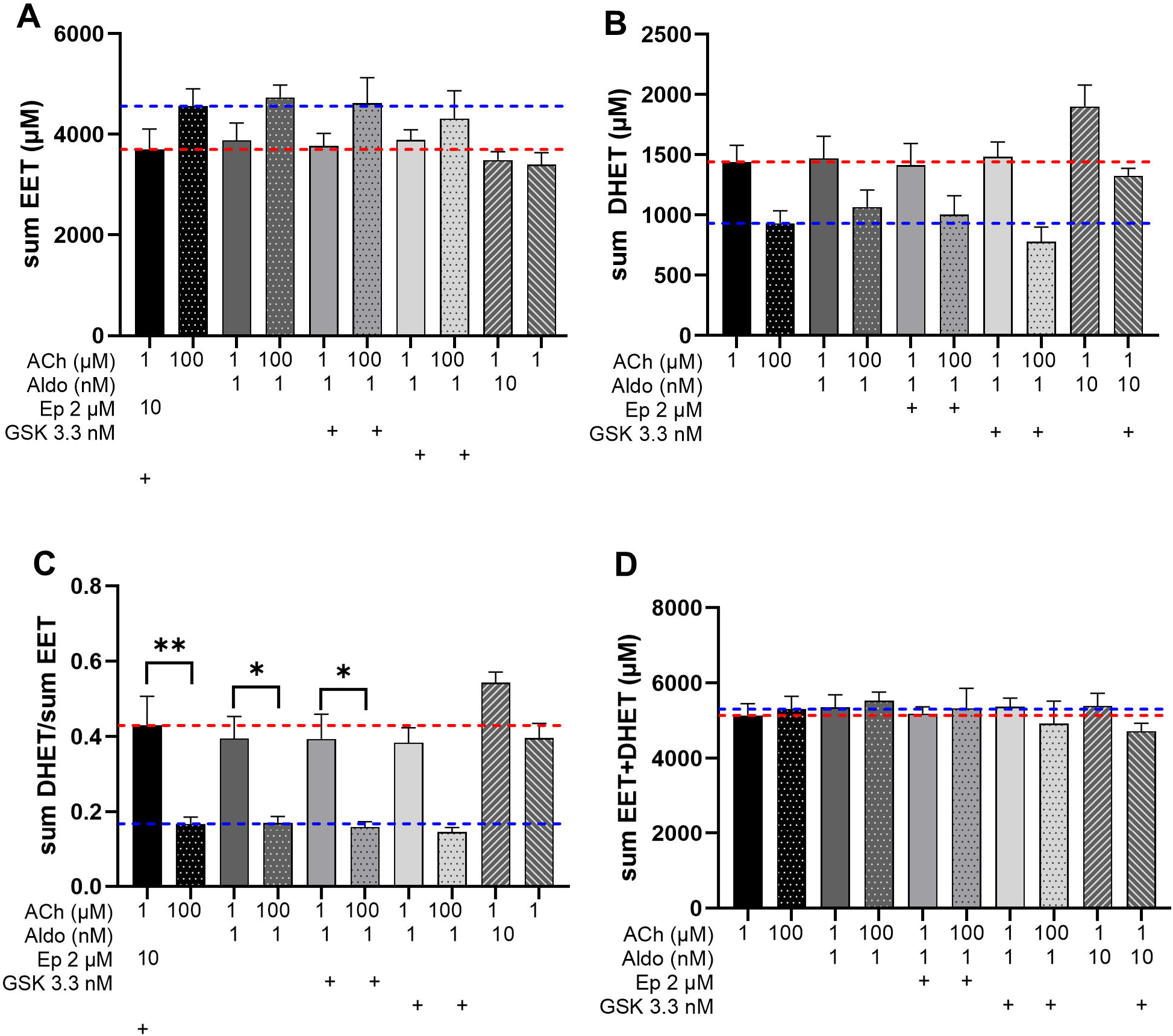
Endothelial cells were synchronized under serum-free conditions, treated with the indicated pharmacological agents in serum-free medium for 48 hours, loaded with 10 µM arachidonic acid and then stimulated either with 1 or 100 µM acetylcholine in HEPES buffer to provoke the release of EETs. Subsequently, supernatants were collected and quantified via HPLC-MS/MS. Dashed bright red line marks the values of control cells stimulated with 1 µM ACh, dashed blue line marks the control cells stimulated with 100 µM ACh. Shown are the sums of all active EETs (5,6; 8,9; 11,12; 14,15 EET, **A**), sum of all inactive EETs (5,6; 8,9; 11,12; 14,15 DHET, **B**), the ratio of the two sums (sum DHET/sumEET, **C**) as a measure of expoxide hydrolase activity and the sum of EET and DHET as a time integral and cumulative measure of EET production as time integral and cumulative measure of total EET production by CYP-epoxigenases (**D**). ACh, acetylcholine; Aldo, aldosterone; Ep, eplerenone; GSK, GSK2256294. N=3-9 per group, *p<0.05, **p<0.01, 1-way ANOVA, Tukey.

Further results showing the treatment response on MR and GR target genes in EC and SMC as well as expression of HSD11B1 and HSD11B2 are provided in the supplement.

We next directly assessed the concentrations of endothelial EETs after exposure to 48 h of aldosterone and cortisol by high pressure liquid chromatography tandem mass spectrometry (HPLC MS/MS). Attention was paid to establishment of the method according to the criteria of the European Medicines Agency[20] (see Supplemental Tables 2 and 3). We used two concentrations of acetylcholine (ACh), 1 and 100 µM. The method was able to quantify all 4 EET regioisomers and their respective DHET epoxide hydrolase products. 14,15 EET and its respective diol were the most abundant regioisomers. ACh-stimulated secretion of EETs was unaltered when compared to cells only treated with cortisol. Likewise, MR antagonism or inhibition of sEH with GSK 2256294 (GSK) parallel to application of aldosterone and cortisol did not resulted in major changes despite that GSK increased 11,12 EET concentrations in ACh 1 µM-stimulated supernatants compared to controls.

Higher concentrations of EETs and lower concentrations of DHETs were measured in the supernatants of cells stimulated with 100 µM *vs*. 1 µM ACh. The sum of EETs and DHETs as a measure of CYP-epoxygenase function did not differ across the two ACh concentrations. Epoxide hydrolase activity was inhibited as a function of ACh concentration, as evidenced by the DHET/EET ratio. In line with this interpretation, GSK precluded the ACh-mediated decrease in 14,15 DHET/EET. Individual concentrations of eicosanoids and derived measures are provided in Supplemental Tables S4 and S5.

EETs reportedly operate indirectly via BK_Ca_-mediated hyperpolarization which leads to a decrease in smooth muscle calcium, relaxation and – eventually – vasodilation[21]. Activation of BK_Ca_ channels in turn is mediated by calcium transients through TRPV4, which are direct targets of EETs[22].

In SMC, TRPV4 expression was unrelated to the presence of aldosterone (Figure 3A). Likewise, no difference in L-type calcium channel expression (CACNA1C coding for Cav1.2) was detected related to the presence or absence of aldosterone with or without cortisol (Figure 3B). Aldosterone alone did not increase CACNA1C transcript levels while cortisol alone reduced Cav1.2 mRNA. Of note, the glucocorticoid receptor antagonist mifepristone (RU486) did not antagonize the cortisol-mediated reduction in Cav1.2 mRNA.

**Figure 3:**
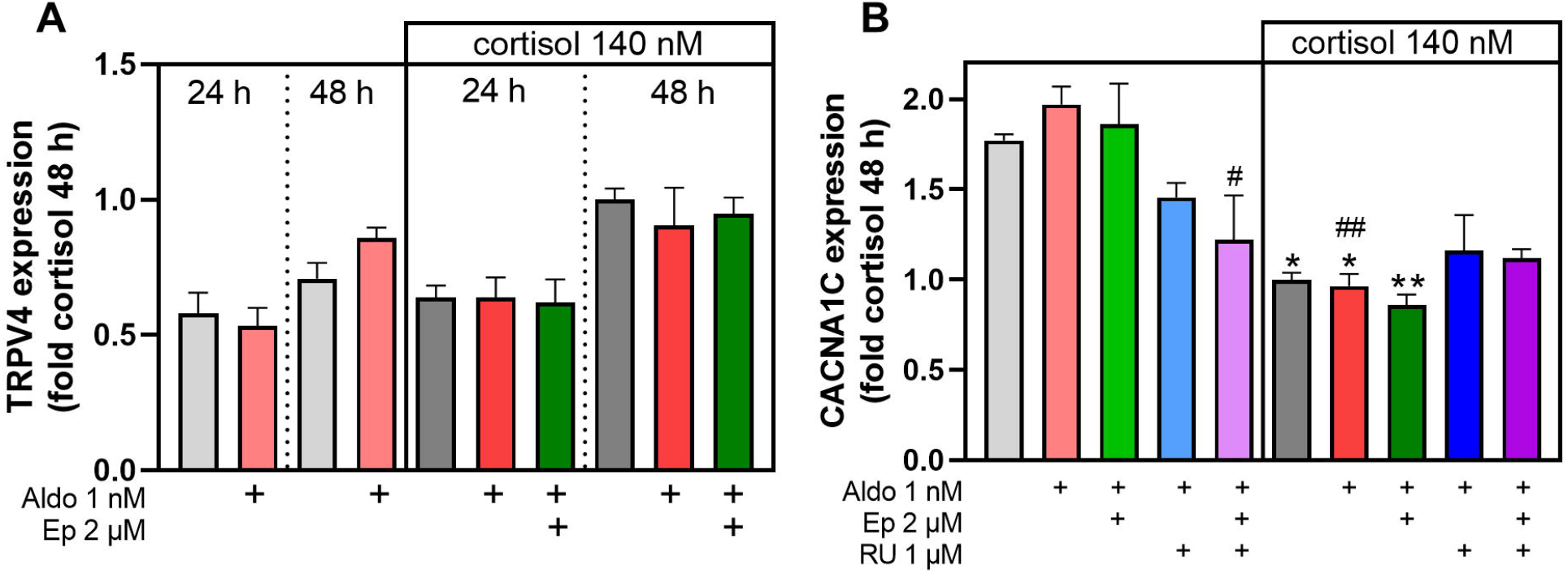
**A:** Effect of duration of exposure to 1 nM of aldosterone and co-stimulation with cortisol on TRPV4 mRNA expression in human coronary artery smooth muscle cells. **B:** Effect of 48 hours exposure to 1 nM of aldosterone and co-stimulation with cortisol on CACNA1C mRNA expression in human coronary artery smooth muscle cells. N=3 per group. Aldo, aldosterone, Ep, eplerenone, RU, mifepristone. *, p<0.05 vs. DMSO; **, p<0.01 vs. DMSO; #, p<0.05 vs. Aldo; ##. P<.0.01 vs. Aldo; 1-Way ANOVA, Tukey.

The expression of BK_Ca_ alpha subunit (KCNMA1) was reduced in SMC treated with aldosterone and cortisol as compared to cortisol alone (Figure 4A). Further examination revealed that the BK_Ca_ message reduction was most pronounced after addition of cortisol. Addition of RU486 to either cortisol or the combined treatment with aldosterone and cortisol substantially increased BK_Ca_ message levels without apparent synergism with eplerenone (Figure 4B). However, despite the reproducible reduction in KCNMA1 mRNA, no effect on protein level could be detected in Western blot (Figure 4C).

**Figure 4:**
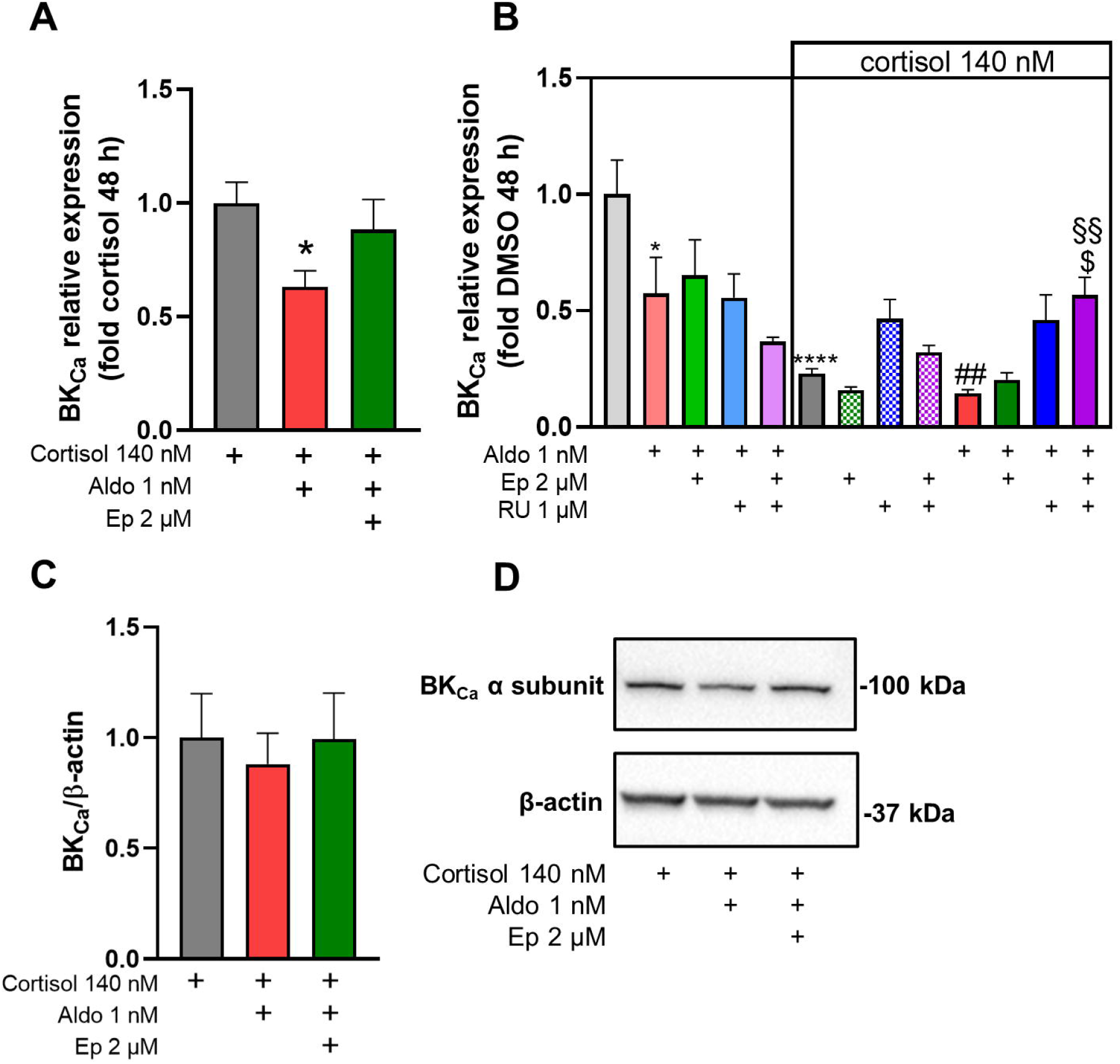
**A:** Effect of exposure to 1 nM of aldosterone and co-stimulation with cortisol on BK_Ca_ alpha subunit mRNA expression in human coronary artery smooth muscle cells. Aldo, aldosterone, Ep, eplerenone. N=6 experiments per group,*p<0.05, Dunnett. **B:** Relative contribution of glucocorticoid receptor stimulation on downregulation of BK_Ca_ alpha subunit mRNA in SMC. Aldo, aldosterone; Ep, eplerenone; RU, RU486 (mifepristone). N=3-6 per group.*p<0.05 vs. control (w/o cortisol), ****,p<0.001 vs. control (w/o cortisol); ##, p<0.01 vs. A 1nM; §§, p<0.01 vs. Aldo+cortisol; $, p<0.05 vs. Aldo+cortisol+Ep. 1-Way-ANOVA, Tukey..**C, D**: Western blot of cortisol, Aldo+cortisol and Aldo+cortisol+Ep-treated SMC for BK_Ca_ alpha subunit and beta-actin. N=4 experiments per group.

14,15 EET-elicited calcium transients in Fura2-loaded SMC were unchanged in cells pre-treated with aldosterone and cortisol for 48 hours when compared to cortisol-treated cells (Figure 5). Application of MRA reduced the calcium transient significantly when compared to the control group.

**Figure 5:**
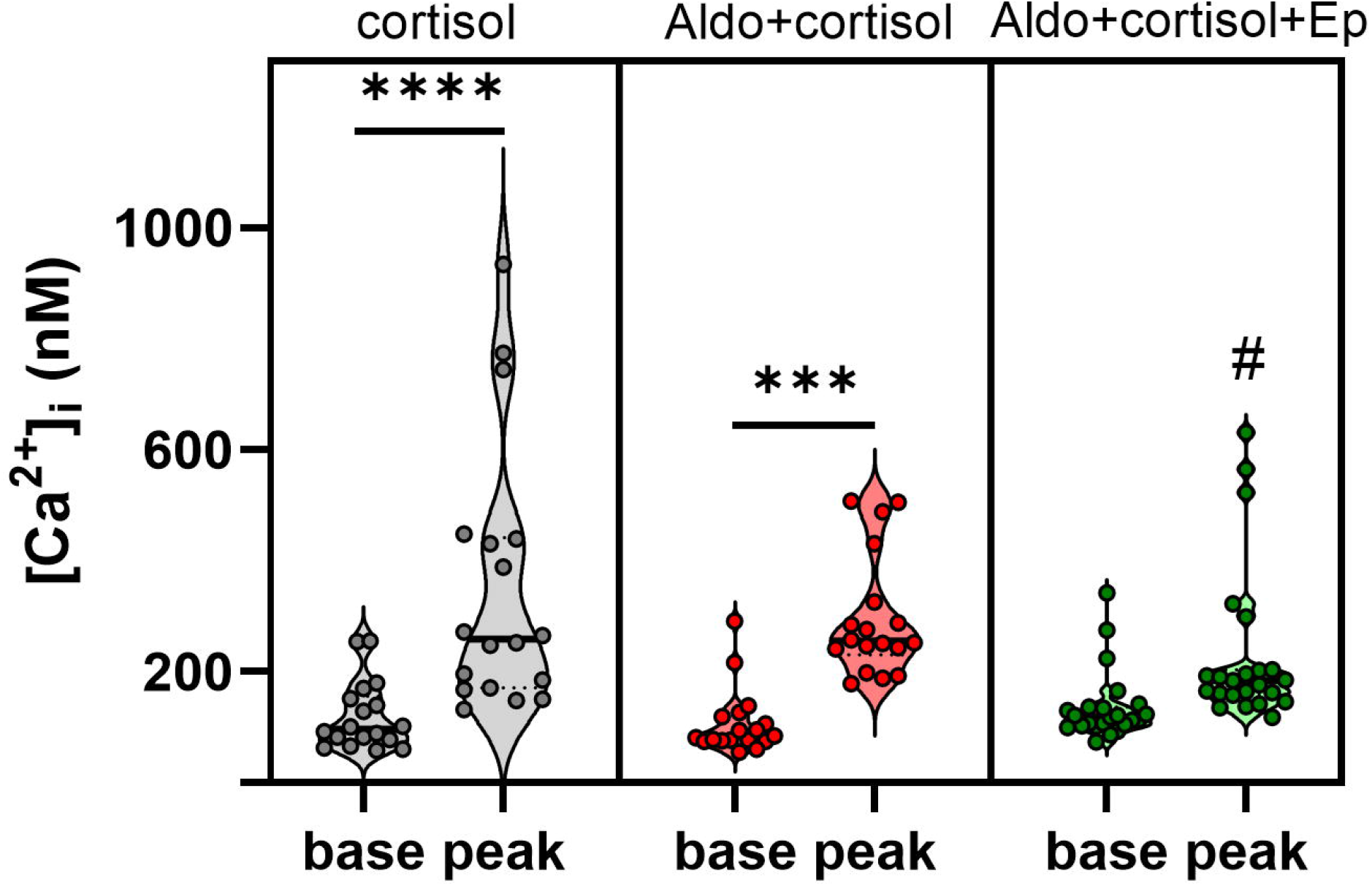
Calcium transient after EET as observed in single smooth muscle cell calcium imaging. Shown are intracellular baseline calcium and peak calcium responses after application of 1 µM 14,15 EET. Aldo, aldosterone; EET, 14,15 epoxyeicostrienoic acid; Ep, eplerenone. ***, p<0.001; ****, p<0.0001;#, p<0.05 vs. peak cortisol (1-Way ANOVA, Tukey), n (cells/total number of experiments)= 17/3 (cortisol), 18/3 (Aldo+cortisol), 23/3 (Aldo+cortisol+Ep).

SMC were submitted to a concentration response curve for 14,15 EET in a plate reader format. A fourth group was included which was treated for 48 hours with 1 µM EET + 3 µM GSK in addition to aldosterone and cortisol. EET and GSK were applied together to correct for SMCs not being able to synthesize EETs themselves. Relative increases in fluorescence were comparable across all three treatment groups until 1 µM EET concentration (Figure 6). At 1 µM, Ep- and EET+GSK-treated cells showed a decline in fluorescence response to EET application. The fluorescence in Aldo-treated cells, however, did not differ to the cortisol-treated control group at any point of the EET concentration-response curve.

**Figure 6:**
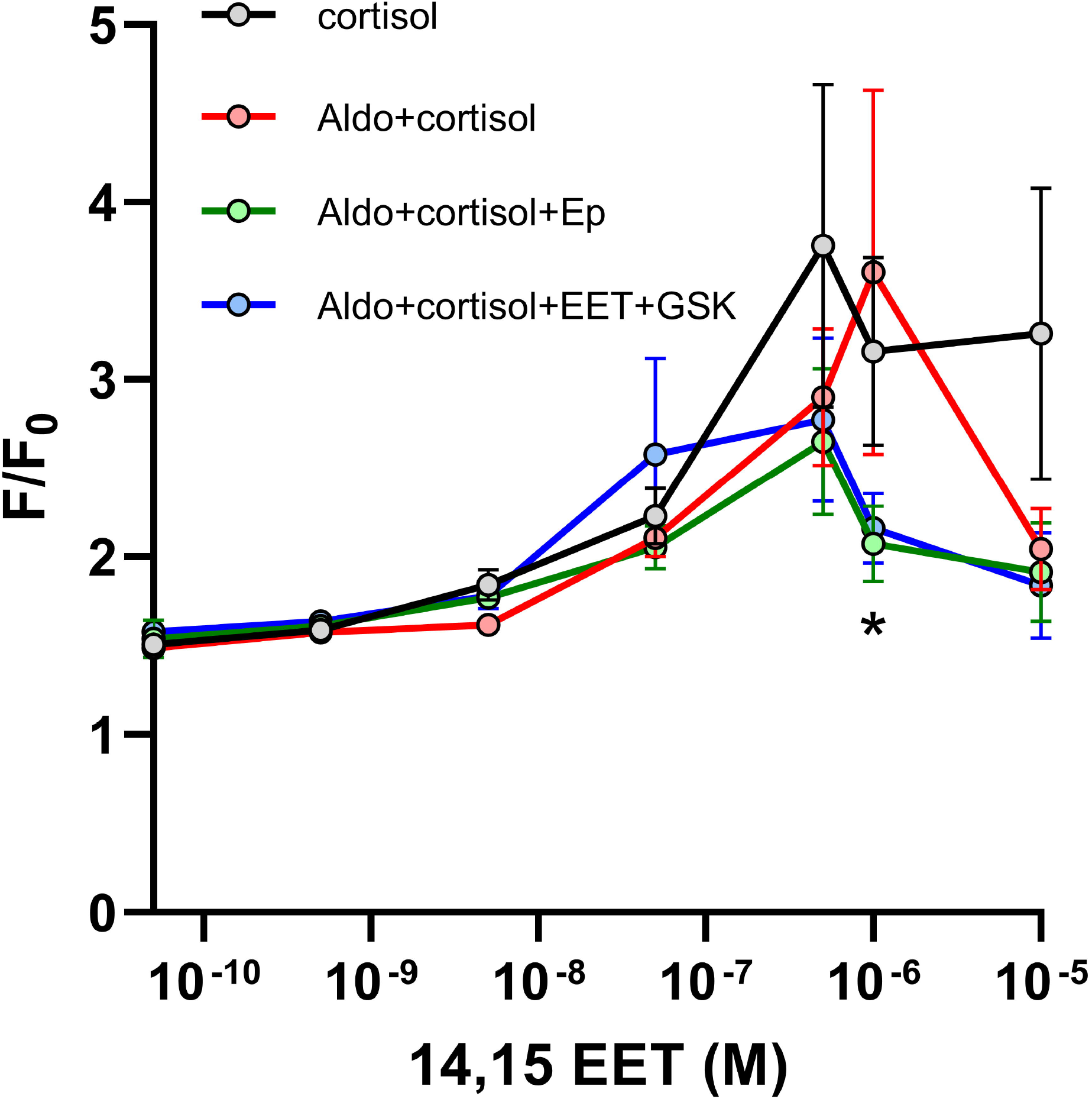
Calcium concentration-response curve for 14,15 EET in fluo4-labelled smooth muscle cells cultured for 48 hours in the presence of the indicated treatments. Aldo, aldosterone; EET, 14,15 epoxyeicostrienoic acid; Ep, eplerenone; GSK, GSK2256294. N=9-21 recordings per concentration. *, p<0.05 (Aldo+cortisol vs. Aldo+cortisol+EET+GSK and Aldo+cortisol vs. Aldo+cortisol+Ep), 2-Way ANOVA, Dunnett.

## Discussion

In the present study we investigated whether concentrations of aldosterone which are typically found in patients with PA impair human coronary artery EC synthetic capacity to release epoxyeicosatrienoic acids (EET). We also addressed whether human coronary artery SMC reactivity to EETs would be hampered by excess of aldosterone. The two cell types were chosen deliberately, since the effects of EETs have been extensively validated in coronary arteries from a variety of species, including bovine[23] and human arteries[24].

Effects of excess of aldosterone on endothelial CYP-epoxygenase expression levels have not been investigated so far. We report that aldosterone essentially does neither alter the expression of CYP2C8 and CYP2J2, nor of epoxide hydrolases which might degrade EETs. Importantly, we were able to corroborate these findings in functional studies: EET release from endothelial cells was not affected by excessive concentrations of aldosterone. In line with our findings, no change in renal expression levels of soluble epoxide hydrolase was detected in a mouse model of DOCA-induced hypertension[25].

We used ACh to trigger endothelial EET release. Two concentrations of ACh yielded a graded endothelial response which was most obvious in the DHET/EET ratio. We therefore concluded that the lack of a difference in stimulated EETs between the treatment groups was not masked by exhaustion of endothelial secretory capacity with maximum ACh concentrations. Consistent with earlier reports, endothelial cells released mostly 14,15 EET and 11,12 EET[26,27], a finding compatible with the preferential synthesis patterns of CYP2C8 and CYP2J2[28,29].

In patients with PA, higher serum levels of 14,15 DHET were detected which was interpreted as increased breakdown of 14,15 EET; however, no ratio of DHET/EET was measured. DHET levels positively correlated with the extent of abdominal aortic calcification[14]. While we also observed numerically higher values of 14,15 DHET in our aldosterone- and cortisol-treated ECs, the differences were not significant. As a caveat, circulating 14,15 DHET and its precursor substance are not necessarily endothelium-derived since other cells such as erythrocytes have been shown to release EETs[30].

The DHET/EET ratio decreased with the higher concentration of ACh. At the same time, the sum of all EETs and DHETs as time integral of EET production remained constant. We therefore conclude that higher concentrations of ACh lead to an inhibition of epoxide hydrolase. It remains to be determined whether in this context the predominant epoxide hydrolase is sEH or mEH. Also, it appears that this study is the first to suggest an ACh concentration-dependent negative regulation of epoxide hydrolase activity.

We detected a minimum 120-fold higher relative expression level of microsomal over soluble epoxide hydrolase in both EC as well as SMC. The negligible effect of GSK in increasing the levels of EETs and decreasing DHET concentrations after stimulation with acetylcholine might be a direct consequence of the low expression of EPHX2. In SMC, we could confirm intact expression levels of both TRPV4 and BK_Ca_ channels as well as Cav1.2 channels, indicating an intact molecular machinery to transduce endothelium-derived EET signals. In line with these findings, calcium responses to EETs were comparable between control cells and aldosterone-treated SMC. We could show synergistic glucocorticoid and mineralocorticoid small repressor effects on BK_Ca_ transcription. The isolated effect of aldosterone, however, did not translate into diminished protein levels as also reflected in unchanged calcium responses to 14,15 EET. We concluded that BK_Ca_ channels are mostly regulated by glucocorticoids and that their transcriptional control is most likely not a primary mechanism of aldosterone-mediated alterations in vascular smooth muscle function. Conflicting data from mouse models on the role of A on BK_Ca_ expression do not allow to integrate our data into a clear picture[31–33].

To the best of our knowledge, this study is the first to suggest glucocorticoid receptor and MR synergism in transcriptional repression of BK_Ca_ channels. It will be intriguing to study these effects further in populations with excessive glucocorticoid secretion (i.e. Cushing disease), co-secretion of gluco-and mineralocorticoids (as often found in primary aldosteronism[34]) or chronic exogenous glucocorticoid administration. The smooth muscle calcium transient to 14,15 EET was unaffected by exposure to aldosterone. MRA antagonism, however, severely blunted the calcium transient to EET. Conceivably, calcium imaging unmasked an inhibitory effect of eplerenone on pannexin 1 hemichannels[35,36] which became obvious at 1 µM EET in both the single cell imaging as well as the plate reader format.

Stimulation with single EET concentrations has previously been shown to increase global smooth muscle intracellular calcium[37]. True concentration-response curves in SMC for cytosolic calcium have not been performed to date.

Of note, while most EET concentrations led to similar responses among all treatments, MR antagonism as well as chronic exposure of aldosterone-treated cells to EET plus sEH inhibitor drastically reduced the calcium response to 1 and 10 µM 14,15 EET. Reduced intracellular calcium should also translate into vasodilation[38]. Therefore, sEH inhibition might constitute a means to increase vasodilation to EETs.

Our study has shown that aldosterone-mediated endothelial dysfunction does not seem to impact on the EET pathway. Experimental trials have shown that soluble epoxide hydrolase inhibition was able to reverse DOCA-induced hypertension and endothelial dysfunction[15]. Thus, although soluble epoxide hydrolase inhibition probably cannot be simply understood as a reversal of the underlying pathophysiological processes, its benefits in conditions of mineralocorticoid excess should be further investigated. Beneficial effects might result from destiffening endothelial cells[39] by reducing ENaC channels[40] which would allow for increased NO release[41]. Another favorable mode of action in mineralocorticoid excess might be EET-mediated attenuation of inflammation[42]. Moreover, EETs have also been described to ameliorate insulin resistance[43]. Since patients with PA frequently show impaired insulin secretion[44], an epoxide hydrolase inhibitor therapy might -again -be of potential additional benefit for patients with PA.

The main limitations of our study consist in its *in vitro* nature precluding any study of heterocellular communication or interaction with systemic factors. In addition, the time scale at which the exposure to aldosterone excess can be studied in primary cell culture systems does not mirror the prolonged exposure in humans.

On the other hand, we believe that a particular point of strength is that we tried to account for both mineralocorticoid and glucocorticoid effects at both steroid receptors, a situation akin to that *in vivo*. Further, our EET quantification underwent an extensive validation protocol according to standards of the European Medicines Agency[20] (Supplemental Tables S2 and S3).

### Perspectives

A substantial proportion of patients with PA still suffers from hypertension even after unilateral adrenalectomy and cure of aldosterone excess [45]. Moreover, medication side effects of MRA hamper therapeutic adherence of patients with bilateral disease. There is, thus, an urgent need for novel pharmacological strategies targeting the downstream effector cells of mineralocorticoid excess. Based on our results, it is therefore tempting to propose that the EET-EDHF axis might be intact in patients with PA. Therapeutic modulation of EET half-lives with epoxide hydrolase inhibitors in conditions of mineralocorticoid excess should, therefore, be investigated in future experiments since the structures responsible for signal transduction in arterial cells – at least in our primary cell culture system – remained intact.

## Acknowledgements

We are indebted to Michaela Höhne and Laura Danner for excellent technical assistance.

