## Supplement for "The epoxyeicosatrienoic pathway is intact in endothelial and smooth muscle cells exposed to aldosterone excess"

**DATA SUPPLEMENT**

**Supplemental Materials and Methods:**

**Cell culture and stimulation**

Human coronary artery endothelial and smooth muscle cells were purchased from Lonza (Basel, CH) together with their specific growth media and growth factors and were cultured in specific growth-medium with supplements according to the manufacturer's recommendations (details see below in section Drugs, buffer compositions, chemicals). All experiments were conducted between passages 5-9 and both cell lines were maintained at 37°C, 5% $CO_2$ and 95% $O_2$ in a humidified incubator. Cells were serum starved over night before treatment in SMBM medium (smooth muscle cells) or EBM-2 (endothelial cells), respectively) as the confluence reached 85-90%. Pharmacological treatment was performed in serum-free medium for an additional 48 hours.

**RNA extraction, reverse transcription and qPCR**

Total RNA was extracted with a cartridge-based Maxwell 16 LEV simply RNA cells kit (Promega, Walldorf, Germany) following to manufacturer's protocol. This protocol includes treatment of the lysates with DNAse I to remove possible contamination with genomic DNA. RNA concentration was measured with a Nanodrop 1000 Spectrophotometer (Thermo Fisher Scientific, Munich, Germany). For cDNA synthesis, RNA was reversed transcribed via the GoScript™ Reverse Transcriptase Kit (Promega,). Quantitative real-time PCR (qRT-PCR) was conducted on the QuantStudio 5 systerm (Thermo Fisher Scientific) using specific Taqman probes (see Supplementary Table S1 below). Relative expression levels of each gene of interest were normalized to the geometric mean of two housekeeping genes (GAPDH and PPIA for SMCs, EIF2B1 and HPRT1 for EC).

| Gene of interest | Taqman probe ID |
|---|---|
| CACNA1C | Hs00167681_m1 |
| CYP2C8 | Hs00946140_g1 |
| CYP2C9 | Hs04260376_m1 |
| CYP2J2 | Hs00559374_m1 |
| EPHX1 | Hs01116806_m1 |
| EPHX2 | Hs00932316_m1 |
| EIF2B1 | Hs00426752_m1 |
| FKBP5 | Hs01561006_m1 |
| GAPDH | Hs00266705_g1 |
| HPRT1 | Hs99999909_m1 |
| HSD11B1 | Hs01547870_m1 |
| HSD11B2 | Hs00388669_m1 |
| KCNMA1 | Hs01119504_m1 |
| PPIA | Hs99999904_m1 |
| SCNN1A | Hs00168906_m1 |
| TRPV4 | Hs01099348_m1 |

**Supplementary Table S1**: Genes of interest and corresponding Taqman probes used in the qPCR experiments.

**Western blot**

SMCs were harvested in RIPA buffer (Thermo Fisher Scientific) supplemented with protease inhibitor cocktail (PIC, Roche, CH) after 48 hours of indicated treatment. Protein concentrations were measured using Pierce BCA Protein Assay Kit (Thermo Fisher Scientific). 20 µg of each lysate were loaded per lane on a 12 % SDS gel (12% Mini-PROTEAN® TGX gels, Bio-Rad, Feldkirchen, Germany) and were transferred onto a PVDF membrane (Bio-Rad US) by Trans-Blot Turbo Transfer system at 5 minutes with 2.5 A current (Bio-Rad). Membranes were incubated with primary antibodies overnight under continuous agitation at 4°C (rabbit anti-

BKCa (1:500, APC-107, Alomone labs, Jerusalem, Israel), mouse anti-β-actin (1:500, Sigma Aldrich, Taufkirchen, Germany)). Membranes were incubated for 1 h with the corresponding secondary antibodies (donkey anti-rabbit HRP conjugate, 1:5000, NA9340V, Amersham, UK; sheep anti-mouse HRP conjugate, 1:5000, NA931V, Amersham) followed by application of detection reagent (Clarity western ECL Substrate, Bio-Rad). Images were acquired with a ChemiDoc XRS+ (Bio-Rad), and band intensities were quantified using ImageJ.

**Fura2-based calcium imaging:**

Cells were grown on glass coverslips with a diameter of 25 mm until 80 % confluence, then serum-starved for 4 hours and treated with 140 nM F, 140 nM F+ 1 nM A or 140 nM F + 1 nM A + 2 µM Ep for 48 hours in serum-free medium. Cells were incubated with 5 µM Fura2-AM in HEPES buffer supplemented with 0.3 % BSA (w/V) for 10 minutes.

After washing with HEPES, coverslips were mounted on a inverted microscope (Olympus IX 71, equipped with a UPlanSApo 20 x /0.85 oil immersion objective and a monochromator (Polychrome V, TILL Photonics, Kaufbeuren, Germany). Signals were recorded with a 14-bit EMCCD camera (iXON 885 K, Andor, Belfast, UK) and digitized using a commercially available software (TILL-VISION, TILL Photonics). Cells were kept under continuous superfusion and stimulated with 1 µM 14,15 EET dissolved in HEPES buffer. As soon as a calcium response occured, the solution was changed to HEPES to avoid overstimulation of the cells. Experiments were conducted at room temperature.

Fura2 F340/F380 ratios (R) were transformed into intracellular calcium concentrations by the following formula[1].

$[Ca^{2+}]_i = K_d*\beta*(R-R_{min})/(R_{max}-R)$

With the experimentally determined values for the setup being

$R_{min} = 0.13$

$R_{max} = 9.722$

$\beta = 16.47$

$K_d = 264$

**Fluo4-based calcium imaging**

SMC were treated as described above in a 96-well plate (Corning® 96 Well CellBIND® Microplates, Sigma Aldrich), loaded with Fluo-4 (5 µM) in HEPES buffer supplemented with 0.1 % pluronic (v/v) (Thermo Fisher Scientific) for 1 hour at 37 °C. Fluo-4 fluorescence signals were acquired with a victor X4 plate reader (PerkinElmer, Walluf, Germany) equipped with injectors to register time-resolved fluorescence after injection. Cells which had undergone the same treatment regimes but which had not been loaded with Fluo4 were included in each run and stimulated with the same EET concentrations to determine non-specific background fluorescence for each EET concentration and treatment. These values were subtracted from the raw fluorescence readings. Background-corrected fluorescence (F) at each timepoint was then normalized to the initial background-corrected fluorescence (F0) to yield F/F0.

**Quantification of EETs and DHETs in endothelial supernatant**

<u>Sample preparation</u>

In a falcon tube, 20 µL of Internal standard solution at 12.5 and 25.0 ng/mL of (±) 11,12-DiHET-d11 and (±) 11,12-EET-d11, respectively (Cayman Chemical, Michigan, USA) and 2.5 mL of cell culture supernatant were extracted with 1000 µL of dichloromethane (DCM) by vortexing for 10 seconds. The bottom layer was transferred to a 2 mL eppendorf tube and the extraction procedure was repeated to ensure the analytes recovery. The DCM extract was dried under vacuum at 40°C, reconstituted in 100 µL of methanol (MeOH) and transferred to a 96 well plate for HPLC-MS/MS analysis. The calibration curve was prepared by spiking standards in HEPES buffer, followed by the same extraction procedure of the samples. Standard from (±) 5,6-EET, (±) 8,9-EET, (±) 11,12-EET, (±) 14,15-EET, (±) 5,6-DHET, (±) 8,9- DHET, (±) 11,12- DHET, and (±) 14,15- DHET (Cayman Chemical, Michigan, USA) were prepared in 7 levels: LOQ, 1, 2.5, 5, 10, 25 and 50 ng/mL.

HPLC-MS/MS

Chromatography was carried out using a 1290 Infinity II high performance liquid chromatography (HPLC) system (Agilent Technologies, Waldbronn, Germany) operating in gradient mode controlled by Analyst software (Version 1.7.0). The chromatographic run was performed using a XBridge® C18 column (2.1 x 150 mm, 3.5 µm) (Waters, Ireland) operated at 40°C. Mobile phase was composed of HPLC-grade water 0.1% formic acid as mobile phase (A), and a mixture of MeOH and acetonitrile (1:1) 0.1% formic acid as mobile phase (B). The flow rate was 1.3 mL.min-1 and the injection volume was 10 µL. Separation was accomplished using a gradient starting with 40% of B kept from 0 to 0.1 min, 70% of B from 0.2 to 3.8 min, 99% of B from 3.9 to 4.4 min, and returning to the initial 40% of B at 4.5 min to 5.5 min.

MS measurements were performed on a QTRAP® 6500+ equipped with Turbo VTM source (AB SCIEX, Toronto, Canada) fully controlled by Analyst software (Version 1.7.0). The instrument was operated with electrospray ionization (ESI) in negative ion mode in multiple reaction monitoring mode (MRM) using scheduled MRM with detection window of 120 sec. The Ion source parameters were: curtain gas (30 psi), IS (-4500 V), gas 1 (50 psi), gas 2 (70 psi), temperature (700°C), collision gas (CAD) (medium). Optimized MS transitions and parameters for individually compound are listed in Supplementary Table S2.

Validation procedure

Evaluation of method performance including limit of detection (LOD), lower limit of quantitation (LLOQ), linearity, accuracy, precision, carryover, extraction efficiency expressed as recovery (R%), and matrix effect expressed as matrix factor (MF) was performed in accordance with the Guideline on bioanalytical method validation from the European Medicines Agency[2]. The results of the validation are listed in the supplementary Table S3.

Linearity and lower limit of quantitation

Quantification was based on internal standard method. Calibration samples were prepared at seven different concentrations. The linear least-square regression was used to determine the mean intercepts, mean slope, and determination coefficients (R2) of the calibration curves.

The LLOQ is defined as the lowest concentration of analyte on the calibration curve with, at least, 5 times the signal of a blank sample and an acceptable accuracy with ±20% of the nominal value and precision ≤20% (CV%) from six replicates for a predefined dynamic range. The LOD is defined as the lowest concentration that gives a reproducible instrument response with S/N ratio ≥3.

Accuracy and precision

Within-run precision and accuracy were determined by analyzing seven replicates of the calibrators for all analytes during a single analytical run. Between –run precision and accuracy were determined by analyzing seven replicates of samples at each level through analytical runs made on five different days.

Extraction recovery

The extraction recovery was calculated in terms of R% = C/B x 100, where C is the analyte peak area of blank matrix (HEPES Buffer) spiked with a reference standard before the extraction and B is the analyte peak area of the matrix spiked with the reference standard after the extraction.

Matrix effect

The matrix effect was calculated in terms of matrix factor (MF), obtained from three concentration levels (low, middle, and high calibrators), each in triplicate. The MF is defined as the analyte peak area ratio of blank matrix spiked with an analyte after extraction to the reference standard containing equivalent amount of the analyte neat sample. If MF is equal to 1, this means that no matrix effect is present; if MF <1, this means that there is ionization suppression, whereas if MF >1, this means that there is ionization enhancement.

Carryover effect

Carryover was investigated by injecting 10 µL of solvent in triplicate immediately after the highest calibration standard and the response was observed at the retention time of the analyte detected. It should not be >20% of the LLOQ response and 5% for the internal standard.

| **Analyte ID** | **Q1** | **Q3** | **RT** | **DP** | **CE** | **CXP** |
|---|---|---|---|---|---|---|
| 5,6 EET | 319,00 | 191,00 | 3,66 | -70 | -16 | -27 |
| 8,9 EET | 319,00 | 154,90 | 3,39 | -85 | -16 | -15 |
| 11,12 EET | 319,00 | 208,00 | 3,18 | -50 | -16 | -19 |
| 14,15 EET | 319,00 | 219,10 | 2,83 | -65 | -14 | -15 |
| 5,6 DHET | 337,00 | 145,00 | 1,92 | -95 | -26 | -19 |
| 8,9 DHET | 337,00 | 127,00 | 1,64 | -65 | -28 | -21 |
| 11,12 DHET | 337,00 | 167,10 | 1,49 | -45 | -24 | -15 |
| 14,15 DHET | 337,00 | 207,00 | 1,13 | -60 | -24 | -11 |

**Supplementary Table S2:** Optimized MS transitions and parameters

| **Compound ID** | **MRM transition** | **Linearity ng.mL$^{-1}$** | **LOQ ng.mL$^{-1}$** | **Precision** | | | | | **R%** | **MF** |
|---|---|---|---|---|---|---|---|---|---|---|
| | | | | **Nominal Concentration ng·mL$^{-1}$** | **Within-run** | | **Between–run** | | | |
| | | | | | **Accuracy (%)** | **CV %** | **Accuracy (%)** | **CV (%)** | | |
| 5,6 EET | 319.0 > 191.0 | 0.50 - 250.00 | 1,000 | 0,500 | 95,07 | 8,25 | 104,94 | 11,37 | 91,00% | 0,60 |
| | | | | 10,000 | 113,00 | 5,19 | 111,54 | 5,36 | 118,00% | 0,64 |
| | | | | 50,000 | 94,70 | 1,43 | 96,88 | 3,20 | 146,00% | 0,50 |
| 8,9 EET | 319.0 > 154.9 | 0.25 - 250.00 | 0,250 | 0,250 | 83,61 | 9,28 | 90,18 | 8,83 | 88,00% | 1,06 |
| | | | | 10,000 | 101,96 | 1,46 | 102,19 | 1,03 | 88,00% | 1,03 |
| | | | | 50,000 | 98,41 | 1,25 | 100,63 | 1,45 | 104,00% | 1,08 |
| 11,12 EET | 319.0 > 208.0 | 0.25 - 250.00 | 0,250 | 0,250 | 83,71 | 10,62 | 95,41 | 10,34 | 99,00% | 1,07 |
| | | | | 10,000 | 101,89 | 2,51 | 101,68 | 1,19 | 88,00% | 1,00 |
| | | | | 50,000 | 98,43 | 0,76 | 100,77 | 1,80 | 95,00% | 0,98 |
| 14,15 EET | 319.0 > 219.1 | 0.25 - 250.00 | 0,250 | 0,250 | 87,63 | 7,20 | 88,99 | 4,88 | 86,00% | 1,01 |
| | | | | 10,000 | 101,27 | 0,77 | 101,96 | 1,13 | 87,00% | 1,03 |
| | | | | 50,000 | 98,76 | 1,80 | 100,86 | 1,57 | 108,00% | 0,99 |
| 5,6 DHET | 337.0 > 145.0 | 0.25 - 250.00 | 0,250 | 0,250 | 82,54 | 2,39 | 95,22 | 10,81 | 81,00% | 1,08 |
| | | | | 10,000 | 101,57 | 0,96 | 101,99 | 0,66 | 76,00% | 1,15 |

| | | | | | | | | | | |
|---|---|---|---|---|---|---|---|---|---|---|
| | | | | 50,000 | 99,83 | 2,70 | 100,37 | 0,36 | 100,00% | 1,66 |
| 8,9 DHET | 337.0 > 127.0 | 0.25 - 250.00 | 0,250 | 0,250 | 94,61 | 1,92 | 93,36 | 11,58 | 93,00% | 1,00 |
| | | | | 10,000 | 99,40 | 1,58 | 102,29 | 1,96 | 87,00% | 1,06 |
| | | | | 50,000 | 100,77 | 1,47 | 100,48 | 0,22 | 95,00% | 1,07 |
| 11,12 DHET | 337.0 > 167.1 | 0.25 - 250.00 | 0,250 | 0,250 | 97,53 | 2,22 | 98,52 | 9,38 | 99,00% | 1,05 |
| | | | | 10,000 | 99,03 | 6,35 | 101,13 | 3,05 | 91,00% | 1,10 |
| | | | | 50,000 | 100,07 | 6,93 | 100,32 | 0,42 | 115,00% | 0,92 |
| 14,15 DHET | 337.0 > 207.0 | 0.25 - 250.00 | 0,250 | 0,250 | 90,70 | 2,76 | 94,99 | 15,26 | 91,00% | 1,02 |
| | | | | 10,000 | 101,11 | 2,52 | 101,46 | 0,66 | 80,00% | 1,08 |
| | | | | 50,000 | 100,27 | 3,23 | 100,49 | 0,40 | 91,00% | 1,04 |

**Supplementary Table S3:** Method validation

**Drugs, chemicals, cells, buffer compositions**

Drugs and chemicals

Acetylcholine: A2661, Sigma Aldrich

Eplerenone: 2397, Tocris, Bristol, UK

Aldosterone: A9477, Sigma Aldrich,

Hydrocortisone: H0888, Sigma Aldrich

14,15 EET: 50651, Cayman Chemical, Michigan, USA

Fluo-4: 6255, Tocris, Bristol, UK

Pluronic™ F-127: P3000MP, Thermo Fisher Scientific

GSK2256294: 2220, Axon Medchem BV, Groningen, The Netherlands

Cells, culture media, growth factors

Endothelial cells: Clonetics™ coronary artery endothelial cells, Lonza, Basel, CH

Endothelial medium: 1xEBM™ 2 Basal Medium（CC-3156, Lonza)

Endothelial growth factors: 1xEGM™-2 MV SingleQuots™ Supplement Pack (CC-4147, Lonza) containing: FBS 25ml, Hydrocortisone 0.2ml, hFGF-B 2ml, VEGF 0.5ml, R3-IGF-1 0.5ml, Ascorbic Acid 0.5ml, hEGF 0.5ml, GA 1000 (=gentamicin + amphotericin B) 0.5ml.

Smooth muscle cells: Clonetics™ coronary artery smooth muscle cells (Lonza)

Smooth muscle medium: 1xSmBM™ Basal Medium (CC-3181, Lonza)

Smooth muscle growth factors: 1xSmGM™-2 SingleQuots™ Supplement Pack (CC-4149, Lonza) containing: Insulin 0.5ml, hFGF-B 1ml, hEGF 0.5ml, GA-1000 0.5ml, FBS 25ml

Buffers

HEPES buffer (in mM): NaCl 140, KCl 5,4, $MgCl_2$ 1, HEPES 10, Glucose 10, $CaCl_2$ x $2H_2O$ 2; pH ad 7,4 with NaOH

**Supplemental results**

Transcriptional activation of SCNN1A (coding for ENaC channel) in human coronary artery smooth muscle cells and endothelial cells was compared after 24 and 48 hours of culture under serum-free conditions in the presence of 1 nM aldosterone or control conditions. A notable, albeit not significant, increase of aldosterone-induced transcript levels was only registered in cells cultured for a period of 48 hours (Supplemental Figure S1).

1 nM of aldosterone elicited reproducible increases in ENaC transcript levels in both cell types, whereas higher concentrations were not associated with higher relative ENaC mRNA concentrations (Supplemental Figure S2). This concentration is well compatible with human pathophysiology as 75 % of patients with PA included in the Munich primary aldosteronism registry display plasma aldosterone concentrations of equal or less than 0.794 nM[3].

Due to the 100-1000 fold plasma concentration abundance of cortisol over aldosterone (Aldo), mineralocorticoid receptors (MR) are usually occupied - but not necessarily stimulated by – cortisol. We therefore chose to culture smooth muscle cells (SMC) and endothelial cells (EC) in the combined presence of 140 nM cortisol and 1 nM Aldo.

In smooth muscle cells (SMC), we observed a statistically significant additive effect in cells treated for 48 hours with cortisol 140 nM and Aldo 1 nM versus cortisol alone with respect to induction of ENaC transcription (Supplemental Figure S3A).

In endothelial cells (EC), treatment with Aldo+cortisol for 48 hours did not increase ENaC expression when compared to cortisol-treated EC. Eplerenone (Ep), a mineralocorticoid receptor (MR) antagonist, increased ENaC expression versus cortisol treatment for 48 hours (Supplemental Figure S3B).

Although there were no treatment-related significant differences within the cell types, SMC depicted robust expression of 11β-hydroxysteroid dehydrogenase type 1 (HSD11B1, mean CT control cells: 27.8 ± 0.1) whereas EC had barely detectable levels (mean CT control cells: 38.3, only 1/3 reactions yielded an amplicon) (Supplemental Figure S4).

For 11β-hydroxysteroid dehydrogenase type 2 (HSD11B2) the differences were less clear-cut: SMC had a mean CT in control cells of 37.13 ± 0.41 and in ECs the value was 40.9 ± 2.4. A direct comparison between EC and SMC expression levels could not be made due to a different set of housekeeping genes used for each cell type.

FKBP5 expression as a glucocorticoid receptor downstream target, however, increased robustly in response to stimulation with cortisol and was completely suppressible by 1 µM RU486 (Supplemental Figure S5).

To assess whether aldosterone impacts the smooth-muscle response to $\alpha_1$ adrenergic stimulation, we conducted a concentration response curve for phenylephrine-induced changes in intracellular calcium. To this end, we employed fluo4-labelled smooth muscle cells, treated for 48 hours with either cortisol, Aldo+cortisol or Aldo+cortisol+Ep and recorded the fluorescence response to different concentrations of phenylephrine in a plate reader format.

We essentially observed no differential effect of Aldo on phenylephrine-induced smooth muscle intracellular calcium increases (Supplemental Figure S7).

**Supplemental figures and tables**

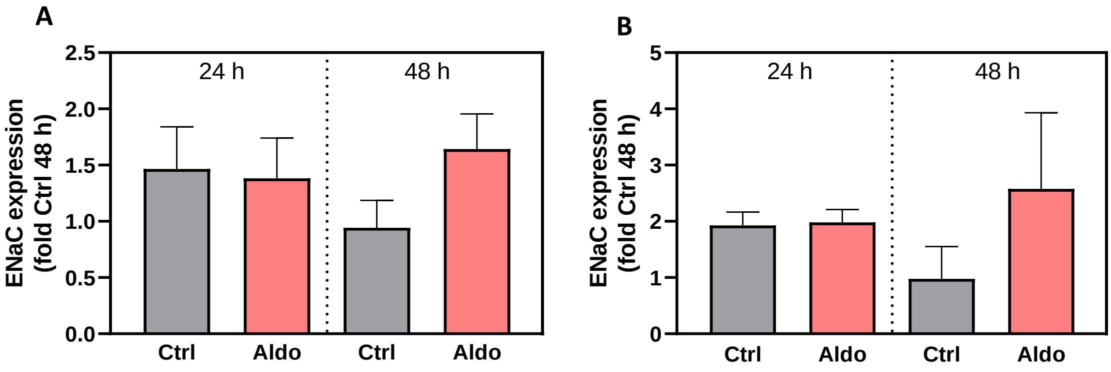


**Figure S1**: Effect of duration of exposure to 1 nM of aldosterone on ENaC mRNA expression in human coronary artery smooth muscle cells (**A**) and endothelial cells (**B**). Each individual ENaC relative abundance was normalized to the mean value of the 48 hours control group. Ctrl, control, Aldo, aldosterone 1 nM. N=3 independent experiments.

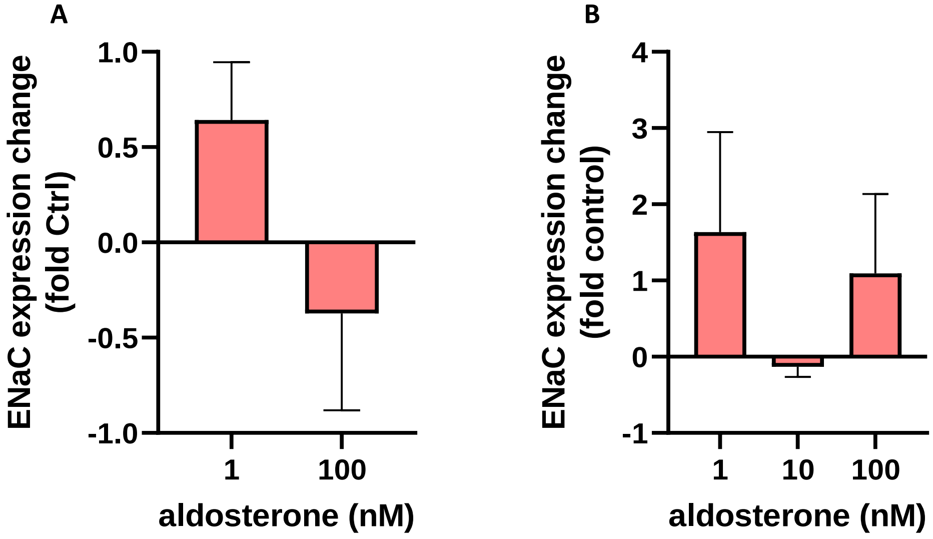


**Supplemental Figure S2**: Effect of aldosterone concentration on ENaC mRNA expression in human coronary artery smooth muscle cells (**A**) and endothelial cells (**B**). ENaC relative abundance at each concentration of aldosterone is expressed as fold change to the appropriate solvent time control. N=3 independent experiments per group.

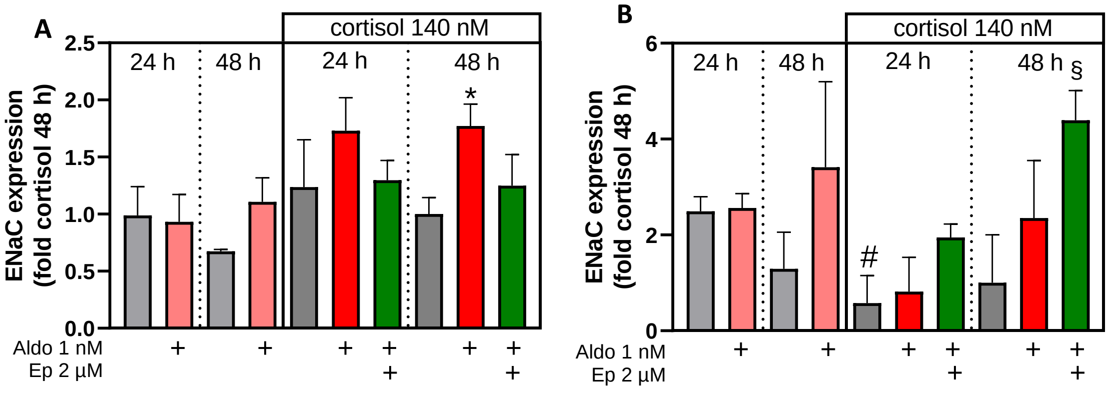


**Figure S3**: Effect of duration of exposure to 1 nM of aldosterone and co-stimulation with cortisol on ENaC mRNA expression in human coronary artery smooth muscle cells (**A**) and endothelial cells (**B**). Aldo, aldosterone, Ep, eplerenone. N=3 in all groups. *p<0.05 vs. cortisol 48 h, t-test. #, p<0.05 vs. 24 h DMSO control, t-test. § p<0.05 vs. cortisol 48 h, t-test.

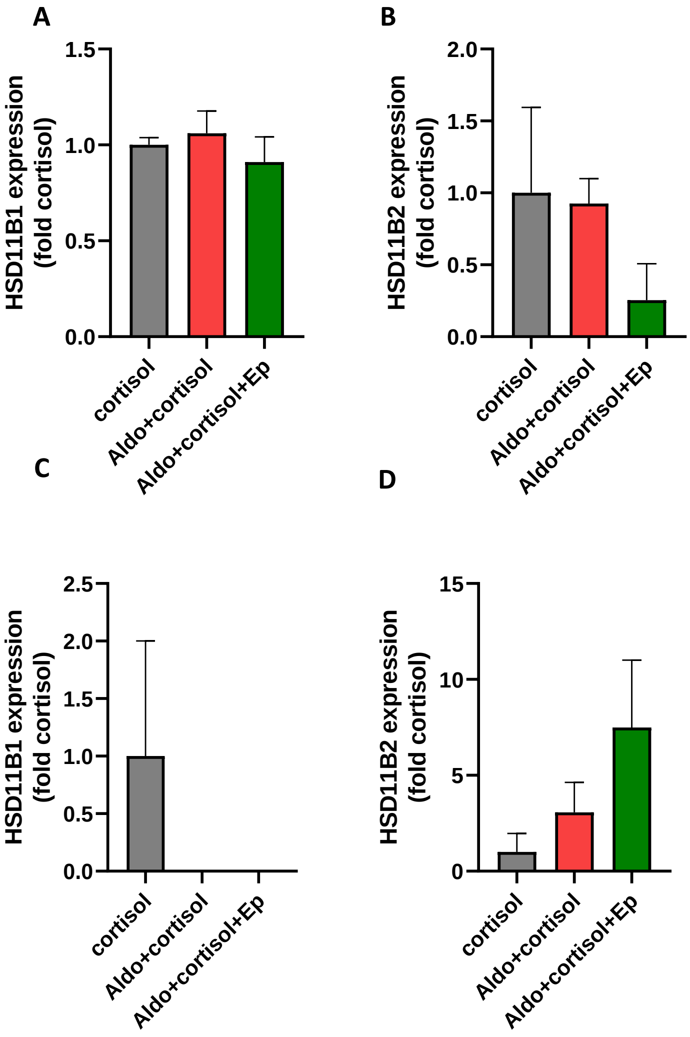


**Supplemental Figure S4**: Effect of exposure to 1 nM of aldosterone and co-stimulation with cortisol on HSD11B1 (**A**) and HSD11B2 (**B**) mRNA expression in human coronary artery smooth muscle cells and endothelial cells (**C**, HSD11B1; **D**, HSD11B2). Aldo, aldosterone, Ep, eplerenone. N=3 independent experiments per group.

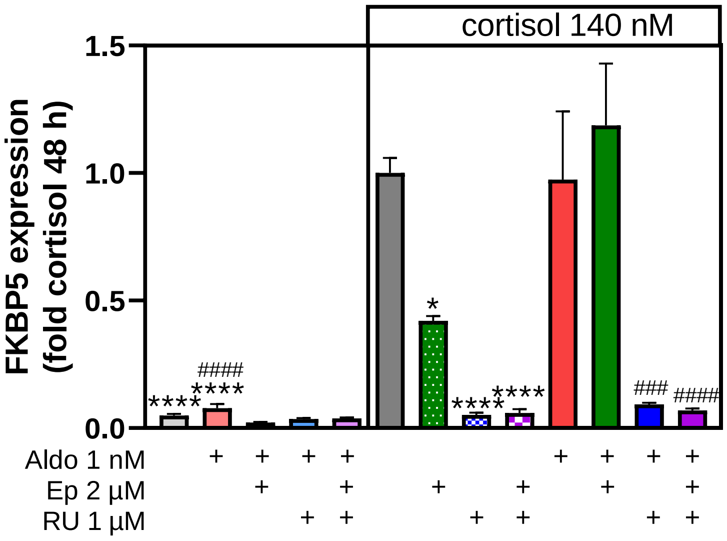

**Supplemental Figure S5**: Effect of exposure of human coronary artery smooth muscle cells to 1 nM of aldosterone and co-stimulation with cortisol on expression of FKBP5 as glucocorticoid receptor downstream target. Aldo, aldosterone, Ep, eplerenone, RU, RU486 (mifepristone);  N=3 independent experiments per group.* $p<0.05$ vs. cortisol; **** $p<0.0001$ vs. cortisol; ### $p<0.001$ vs. Aldo+cortisol; #### $p<0.0001$ vs. Aldo+cortisol, 1-Way-ANOVA, Tukey. n=3 experiments per group.

### Fura2-based calcium imaging

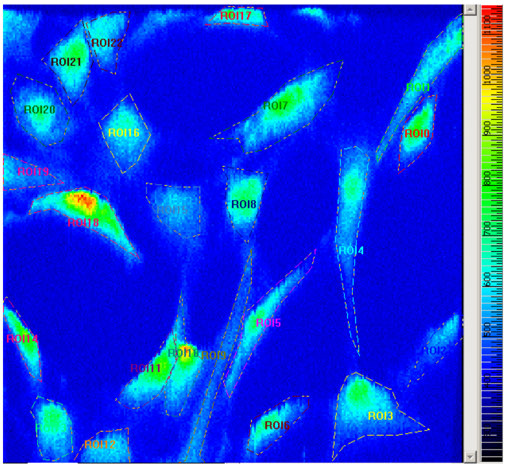


**Supplemental Figure S6:** Example ROIs of SMC treated for 48 hours with aldosterone and cortisol (image acquired before stimulation).

### Fluo4-based calcium imaging

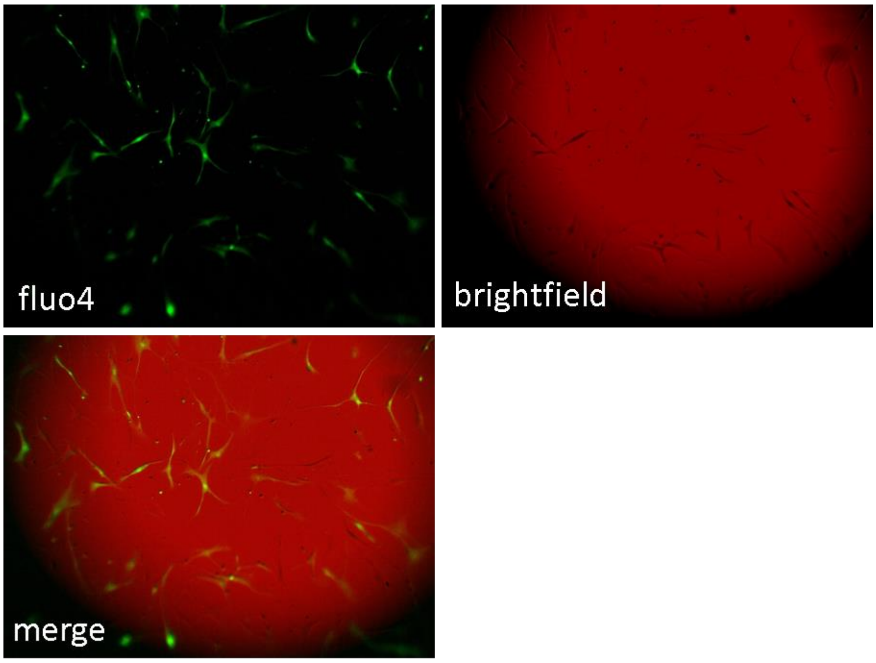


**Supplemental Figure S7:** SMC labelled with fluo4 were imaged with fluorescence microscopy at 5x magnification directly in the 96-well plate before an experiment. Top left: fluo4-specific wavelength. Top right: brightfield image; bottom left: merged image.

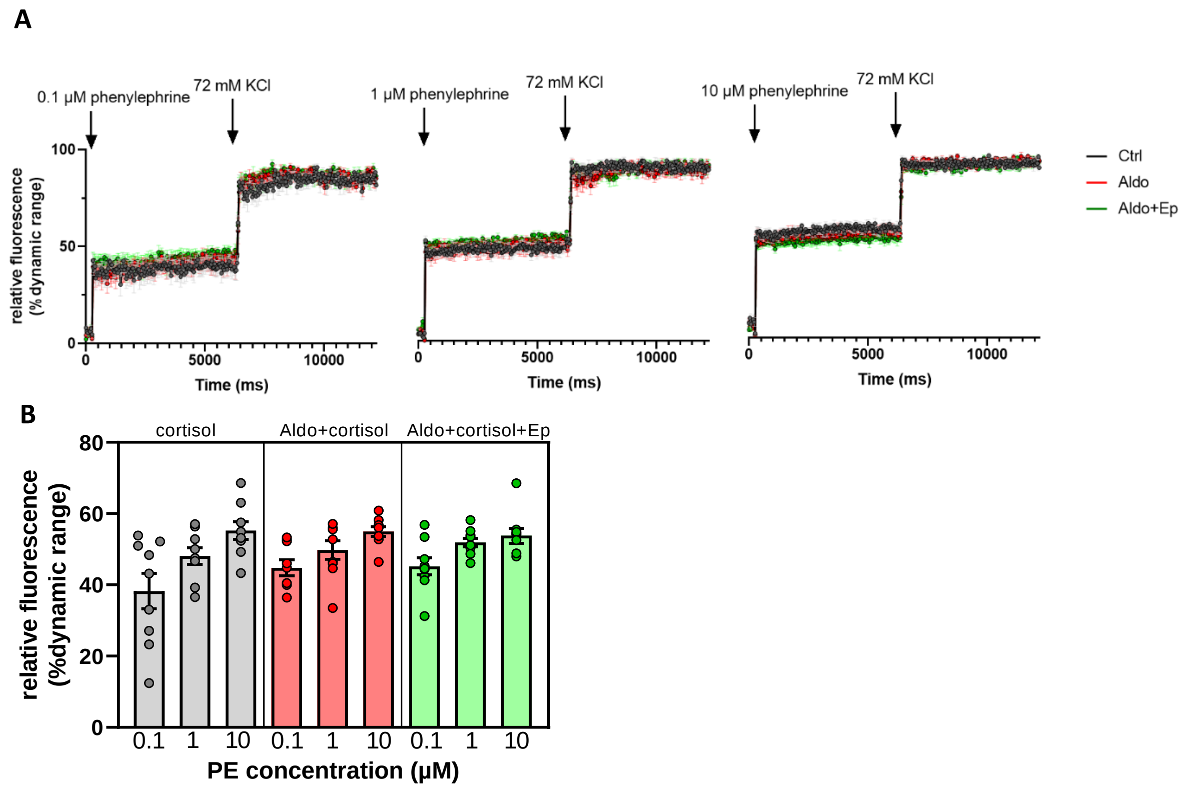


**Supplemental Figure S8:** Time-resolved calcium imaging in SMC exposed to different concentrations of phenylephrine (PE). At the end of each concentration, 72 mM KCl were applied to elicit a maximal calcium response. **A**: summary of 9 individual experiments (mean ± SEM). **B**: PE concentration response curve for the 3 individual treatments. Aldo, aldosterone, Ep, eplerenone; PE, phenylephrine.

| Concentrations of EET and DHET after stimulation of endothelial cells with 1 µM acetylcholine | | | | | | |
|---|---|---|---|---|---|---|
| **Treatment (n)** | **cortisol** (6) | Aldo1+ cortisol (6) | Aldo1+ cortisol+Ep (6) | Aldo1+ cortisol +GSK (3) | Aldo10+ cortisol (3) | Aldo10+ cortisol +GSK (3) |
| **CYP epoxides (µmol/L)** | | | | | | |
| 5,6 EET | 895,59 (195.28) | 813,75 (174.80) | 862,54 (196.68) | 463,71 (114.39) | 400,14 (20.63) | 398,15 (398.15) |
| 8,9 EET | 623.81 (75.24) | 678.44 (88.46) | 636.24 (37.50) | 628.16 (52.73) | 554.35 (32.38) | 574.49 (72.98) |
| 11,12 EET | 932.66 (74.05) | 1048.26 (68.98) | 946.66 (32.29) | **1214.07*** **(13.55)** | 1066.93 (23.41) | 1001.88 (68.28) |
| 14,15 EET | 1249.01 (77.63) | 1344.54 (107.90) | 1325.72 (45.51) | 1585.00 (50.51) | 1472.55 (146.82) | 1422.04 (46.57) |
| Sum EETs | 3701.07 (407.61) | 3884.98 (343.27) | 3771,06 (249.70) | 3890.93 (201.68) | 3493.96 (165.22) | 3396.56 (238.71) |
| **Epoxide hydrolase diols (µmol/L)** | | | | | | |
| 5,6 DHET | 123.47 (21.06) | 124.35 (30.70) | 128.11 (30.67) | 197.00 (15.39) | 172.98 (30.91) | 150.81 (10.46) |
| 8,9 DHET | 39.20 (5.33) | 39.07 (3.99) | 39.88 (5.28) | 55.48 (6.14) | 46.15 (7.58) | 39.41 (2.99) |
| 11,12 DHET | 257.89 (48.71) | 236.34 (38.78) | 238.29 (37.92) | 322.55 (26.59) | 323.80 (37.74) | 272.23 (4.63) |
| 14,15 DHET | 1022.22 (76.35) | 1071.35 (114.62) | 1010.31 (110.12) | 908.70 (78.48) | 1360.87 (102.84) | 864.86 (55.20) |
| Sum DHETs | 1442.79 (137.58) | 1471.11 (183.89) | 1416.58 (177.51) | 1483.72 (122.84) | 1903.80 (176.12) | 1327.31 (62.12) |
| **CYP epoxygenase activity (µmol/L)** | | | | | | |
| Sum EETs+DHETs | 5143.86 (312.95) | 5356.09 (331.30) | 5187.65 (183.90) | 5374.66 (228.38) | 5397.76 (335.47) | 4723.87 (212.18) |
| **Epoxide hydrolase activity (ratio)** | | | | | | |
| 5,6 DHET/EET | 0.20 (0.07) | 0.23 (0.09) | 0.25 (0.10) | 0.49 (0.15) | 0.44 (0.09) | 0.39 (0.05) |
| 8,9 DHET/EET | 0.07 (0.01) | 0.06 (0.01) | 0.07 (0.01) | 0.09 (0.02) | 0.08 (0.01) | 0.07 (0.01) |
| 11,12 DHET/EET | 0.30 (0.07) | 0.22 (0.03) | 0.25 (0.04) | 0.27 (0.02) | 0.30 (0.03) | 0.27 (0.02) |

| | | | | | | |
|---|---|---|---|---|---|---|
| 14,15 DHET/EET | 0.85 (0.10) | 0.81 (0.07) | 0.76 (0.07) | 0.57 (0.05) | 0.93 (0.03) | 0.61 (0.05) |
| Sum DHETs/sum EETs | 0.43 (0.08) | 0.40 (0.06) | 0.39 (0.07) | 0.38 (0.04) | 0.54 (0.03) | 0.40 (0.04) |

**Supplemental Table S4:** EET and DHET in EC supernatant after stimulation with **1 µM** acetylcholine. Values as mean (SEM). Aldo1, aldosterone 1 nM; DHET, dihydroxyeicosatrienoic acid; EC, endothelial cells; EET, epoxyeicosatreienoic acid; Ep, eplerenone 2 µM; GSK, GSK 2256294 3.3 nM; Aldo10, aldosterone 10 nM. *, $p<0.05$ vs. cortisol; 1-Way ANOVA, Dunnett.

| Concentrations of EET and DHET after stimulation of endothelial cells with 100 µM acetylcholine | | | | |
|---|---|---|---|---|
| **Treatment (n)** | **cortisol (9)** | **Aldo1+cortisol (9)** | **Aldo1+cortisol+ Ep (9)** | **Aldo1+cortisol+ GSK (8)** |
| **CYP epoxides (µmol/L)** | | | | |
| 5,6 EET (n=6) | 1082.37 (185.06) | 1092.35 (148.07) | 1051.58 (188.96) | 1087.36 (207.49) |
| 8,9 EET | 819.42 (73.23) | 874.76 (51.56) | 847.54 (82.85) | 836.27 (112.10) |
| 11,12 EET | 1123.84 (99.31) | 1276.18 (60.67) | 1230.42 (97.43) | 1094.37 (117.02) |
| 14,15 EET | 1176.05 (43.30) | 1336.27 (56.80) | 1332.66 (80.86) | 1118.44 (70.95) |
| Sum EETs (n=6) | 4562.05 (347.24) | 4739.08 (244.50) | 4622.17 (507.37) | 4315.74 (553.67) |
| **Epoxide hydrolase diols (µmol/L)** | | | | |
| 5,6 DHET | 144.97 (20.46) | 165.29 (25.81) | 154.59 (25.00) | 120.58 (16.76) |
| 8,9 DHET | 32.89 (4.47) | 38.36 (3.89) | 35.14 (2.89) | 31.82 (3.47) |
| 11,12 DHET | 129.51 (18.52) | 148.72 (19.02) | 136.82 (17.29) | 118.04 (14.55) |
| 14,15 DHET | 625.44 (64.82) | 713.11 (97.13) | 678.06 (114.61) | 506.72 (91.64) |
| Sum DHETs | 932.80 (103.09) | 1065.49 (144.39) | 1004.60 (155.85) | 777.16 (124.26) |
| **CYP epoxygenase activity (µmol/L)** | | | | |
| Sum EETs+DHETs (n=6) | 5306.73 (343.00) | 5533.17 (231.04) | 5330.06 (532.53) | 4920.29 (602.65) |
| **Epoxide hydrolase activity (ratio)** | | | | |
| 5,6 DHET/EET (n=6) | 0.12 (0.03) | 0.12 (0.02) | 0.12 (0.02) | 0.10 (0.02) |
| 8,9 DHET/EET | 0.05 (0.01) | 0.05 (0.01) | 0.05 (0.01) | 0.04 (0.01) |
| 11,12 DHET/EET | 0.13 (0.03) | 0.12 (0.02) | 0.12 (0.02) | 0.12 (0.02) |
| 14,15 DHET/EET | 0.55 (0.08) | 0.54 (0.07) | 0.51 (0.08) | 0.45 (0.08) |
| Sum DHETs/sum EETs (n=6) | 0.17 (0.02) | 0.17 (0.02) | 0.16 (0.01) | 0.14 (0.01) |

**Supplemental Table S5:** EET and DHET in EC supernatant after stimulation with **100 µM** acetylcholine. Values as mean (SEM). Aldo1, aldosterone 1 nM; DHET, dihydroxyeicosatrienoic acid; EC, endothelial cells; EET, epoxyeicosatreienoic acid; Ep, eplerenone 2 µM; GSK, GSK 2256294 3.3 nM. 1-Way ANOVA, Dunnett.
